# Simultaneous Prediction of Functional States and Types of *cis*-regulatory Modules Reveals Their Prevalent Dual Uses as Enhancers and Silencers

**DOI:** 10.1101/2024.05.07.592879

**Authors:** Sisi Yuan, Pengyu Ni, Zhengchang Su

**Affiliations:** Department of Bioinformatics and Genomics, the University of North Carolina at Charlotte, Charlotte, NC, 28223, USA

**Keywords:** cis-regulatory modules, enhancers, silencers, functional state, dual functional CRMs

## Abstract

Our understanding of the precise locations of *cis*-regulatory elements (CRMs) in the genomes, as well as their functional types (enhancer or silencer), states (active or inactive) and target genes in various cell/tissue types of organisms remains limited, despite recent progresses. To address these challenges, we have recently developed a two-step strategy that first predicts a more complete map of CRMs in the genome, and then predicts the functional states of the CRMs. However, our initial approach lacked the ability to differentiate between the functional types of CRMs. Therefore, we utilized distinct features to simultaneously predict the functional types and states of the CRMs. Applying our method to 107 cell/tissue types with the minimum of required data available, we predicted 868,948 (73.8%) of the CRMs to be active as enhancers or silencers in at least one of these cell/tissue types. In 56 cell/tissue types with required data available for both enhancers and silencers, we predicted that 117,646 (14.8%) and 227,211 (28.6%) CRMs only functioned as enhancers (enhancer-predominant) and silencers (silencer-predominant), respectively, while 83,985 (10.6%) functioned both as enhancers and silencers (dual functional). Thus, both dual functional CRMs and silencers might be more prevalent than previously assumed. Most dual functional CRMs function either as enhancers or silencers in different cell/tissue types (Type I), while some have dual functions regulating different genes in the same cell/tissue types (Type II). Different types of CRMs display different lengths and TFBS densities, reflecting the complexity of their functions. Our two-step approach can accurately predict the functional types and states of CRMs using data of only five epigenetic marks in a cell/tissue type.

**Author Summary:** CRMs function as enhancers and/or silencers to promote and repress, respectively, the transcription of genes in a spatiotemporal manner, thereby playing critical roles in virtually all biological processes. However, despite recent progress, the understanding of CRMs remains limited. Most existing methods are aimed to simultaneously predict the locations and functional states of enhancers in a given cell/tissue type, however, the accuracy of these one-step methods is low. We have recently developed a two-step strategy that first predicts locations of CRMs in the genome, and then predicts their functional states as enhancers in cell/tissue types with high accuracy. However, our initial approach was unable to differentiate between enhancers and silencers. Therefore, in this study, we employ two machine-learning models, so that we can simultaneously predict the functional states and types of our previously predicted 1.2M CRMs in various cell/tissue types. Applying the method to cell/tissue types with the data available, we categorize the CRMs into four types with distinct properties reflecting their functional complexity. Our results indicate that silencers and dual functional CRMs might be more prevalent than previously assumed. The precise prediction of CRM types and states provides opportunities to pinpoint their target genes, thus opening new avenues for research.

## Introduction

Approximately 99.5% of genomes among human individuals are identical at the single nucleotide level[1], suggesting that the remaining 0.5% of genetic variation, mostly located in non-coding regions[2, 3], might account for the diverse phenotypes within the human populations. Consistently, genome-wide association studies (GWAS) have unveiled that nearly 90% of single nucleotide polymorphisms (SNPs) associated with complex diseases or phenotypes reside in non-coding regions[4]. This underscores that disparities in phenotypes and disease susceptibilities across individuals primarily stem from variation within non-coding regions, particularly those that disrupt transcription factor (TF) binding sites (TFBSs) in CRMs located predominantly in non-coding regions[5–8]. CRMs, classified as enhancers, silencers, promoters and insulators based on their effects and roles in transcriptional regulation, often exhibit on-off switches in functional states across different cell/tissue types[9, 10]. Particularly, specific TFs binding to cognate binding sites in an enhancer or a silencer can facilitate or prevent, respectively, the recruitment of RNA polymerase to the promoter of the target gene, thereby upregulating or downregulating transcription of the gene, respectively[11, 12]. Some TFs such as CTCF bind to their cognate sites within insulators to establish functional domain boundaries that confine the regulatory actions of the CRMs within the designated topologically associating domain (TAD)[13]. Epigenetic modifications of CRMs within diverse cell types can influence the accessibility and binding affinity of TFs to their cognate binding sites. Consequently, this dynamic interplay among TFs, TFBSs, RNA polymerase and epigenetic modification systems contributes to distinct spatiotemporal expression patterns of genes in various biological processes across different cell/tissue types[14, 15].

CRMs have long been characterized by using low throughput laborious molecular biology methods. However, recent advancements in a plethora of omics techniques have revolutionized our capacity to investigate CRMs. These techniques encompass: 1) chromatin immunoprecipitation sequencing (ChIP-seq) for identifying TFBSs[16–18] and regions modified by histone marks[19] in the genome; 2) DNase I hypersensitive sites sequencing (DNase-seq)[20–22] and assay for transposase-accessible chromatin using sequencing (ATAC-seq)[23] for probing chromatin accessibility (CA) of genome regions; 3) Hi-C technology for measuring physical proximity between genomic loci in the nucleus[24, 25]; and 4) RNA-seq for quantifying transcriptomes in cells/tissues[26]. The wide adaptations of these techniques have resulted in vast volumes of data, originating from large consortia as well as individual laboratories worldwide[27–34]. This wealth of data presents an unparalleled opportunity to reliably predict the location of CRMs in genomes, along with their functional states (on/active or off/inactive), types (predominantly enhancers or silencers), and target genes across diverse cell/tissue types[35, 36]. Most existing methods attempted to simultaneously predict locations and functional states of enhancers in a given cell/tissue type by integrating multiple epigenetic marks including CA and various histone modifications[37–41]. Although conceptually appealing, these one-step methods are limited for their high false discovery rates (FDRs)[42–48]. This is due to the fact that the presence of CA and histone marks, while informative, is not exclusively indicative for enhancers and their functional states, as these marks are also present in non-CRM sequences[45, 46, 49].

On the other hand, it has been shown that TF binding data are more informative for identifying loci of CRMs than CA and histone modification data[45–49]. Furthermore, it has been established that accurate anchoring of a CRM’s location through the binding of key TFs renders epigenetic marks on the CRM a reliable predictor of its functional states[45–49]. In light of these findings, we have recently introduced a two-step approach to predict CRMs and their functional states sequentially[42, 50]. Firstly, we predict an accurate and more complete map of CRMs in the genome using TF binding data. Secondly, we predict the functional states of all the predicted CRMs in any given cell/tissue type of the organism using few epigenetic marks. For the first step, we have developed the dePCRM2 algorithm[42], which predicts loci of CRMs by integrating putative TF binding motifs identified in a large number of diverse TF ChIP-seq datasets using our ultra-fast motif finder ProSampler[51, 52]. dePCRM2 is able to effectively segregate genomic regions covered by TF binding peaks into two exclusive sets: the CRM candidates and the non-CRMs[42]. While dePCRM2 can predict a CRM’s functional state in a specific cell/tissue type based on its overlaps with TF binding peaks available in the very cell/tissue type, this predictive capacity is often limited due to the scarcity of available TF binding datasets in most cell/tissue types. Therefore, similar to the candidate *cis*-regulatory elements (cCREs) identified recently by the ENCODE project[53], our predicted CRMs are generally cell/tissue type agnostic. For the second step, we have developed a machine learning model that can accurately predict the functional states of all the predicted CRMs as enhancers in diverse human and mouse cell/tissue types using only four epigenetic marks as features[50]. This two-step approach significantly surpasses existing one-step methods in terms of sensitivity and specificity for predicting active enhancers in various cell/tissue types[50].

However, recent investigations have found that silencers are more prevalent than initially believed[54–56], and that an active enhancer in a cellular context could be an active silencer in another cellular context[9, 56]. Thus, it is interesting to also predict functional states of CRMs as silencers. Indeed, a few diverse computational tools have been developed to predict active silencers in specific cell/tissue types[55]. However, as in the case of enhancers, these methods attempted to simultaneously predict the locations and functional states of silencers using epigenetic marks on candidate DNA segments[57–59]. For example, a correlation-based method correlated putative active silencer mark (e.g., H3K27me3 and DNaseI hypersensitive site (DHS)) signals with the expression levels of neighboring genes across different cell/tissue types[57]. A support vector machine (SVM) model was then trained using sequence features as well as features derived from the aforementioned correlation-based method to predict silencers[57]. Additionally, a simple subtractive approach (SSA) excluded genome regions with enhancer chromatin signatures to be putative silencers and a gapped *k-mer* SVM (gkmSVM) was trained on massively parallel reporter assay (MPRA) data and sequence patterns to predict silencers[59]. However, the accuracy of these one-step methods is quite low (see later), due to similar reasons for predicting enhancers and their functional states using one-step methods.

Building upon the success of our two-step strategy in predicting enhancers and their functional states in cell/tissue types, we now extend our machine learning model to simultaneously predict the functional types (enhancer or silencer) and states (on/active or off/inactive) in various cell/tissue types in a genome-wide fashion. Our methods achieve an area under the receiver operator characteristic curve (AUROC) > 0.96 and show superior performance to state-of-the-art methods. Using the tools, we predicted functional types and states of 1.2M CRMs in 107 human cell/tissue types. Our results indicate that silencers and dual functional CRMs are more prevalent than previously thought and that various types of CRMs display distinct properties in terms of their lengths and TFBS densities, reflecting their functional complexity.

## Results

### Functional states of CRMs as silencers and enhancers can be accurately predicted using three epigenetic marks

We have previously developed a logistic regression (LR) model to predict functional states in a cell/tissue type of our 1.2 M predicted CRMs in the human genome[42] using signals of few epigenetic marks on the CRMs as features that are more or less associated with active enhancers. We employed a similar LR model (Figure 1A) to predict functional states of the CRMs as silencers in a cell/tissue type using three epigenetic marks (CA, H3K9me3 and H3K27me3) on the CRMs as features. We pooled positive and negative silencer sets compiled in each of 40 of the 67 human cell/tissue types with the required data available (Materials and Methods), resulting in a positive set containing a total 256,766 positive silencers and a negative set with the same number of negative control sequences. As shown in the UpSet plot in Figure 1B, H3K27me3 peaks pooled from the 40 cell/tissue types have the highest coverage of the human genome, followed by CA and H3K9me3 peaks, and around 100 Mb of the genome are covered by the peaks of all the three marks. To evaluate the ability of these three marks to predict the functional states of the CRMs as silencers in cell/tissue types, we trained and evaluated the seven models using all the seven possible combinations of one, two and three of the three marks as features by 10-fold cross validation (Figure 1B). Of the three models using only one mark, model 2 using CA had the highest median AUROC of 0.948, followed by model 1 using H3K27me3 (median AUROC=0.826) and model 3 using H3K9me3 (median AUROC=0.685). Thus, CA alone has quite high prediction accuracy, while H3K27me3 alone and particularly, H3K9me3 alone have only intermediate prediction accuracy. Of the three models using two marks, model 4 using CA and H3K27me3 (CA&H3K27me3) obtained the highest median AUROC of 0.960, followed by model 6 (CA&H3K9me3, median AUROC=0.951) and model 5 (H3K9me3&H3K27me3, median AUROC= 0.919). Model 7 using all the three marks (CA&H3K9me3&H3K27me3) achieved the highest median AUROC of 0.962 (Figures 1B, 1C), which is significantly higher than the other six models (p value < 0.01, Mann-Whitney U test). Consistently, CA in model 7 had a much higher weight (102.7) than H3K27me3 (16.5) and H3K9me3 (10.9) (Figure 1D). We thus selected model 7 as our silencer functional state predictor in the subsequent predictions. The numbers of predicted active silencers in these 40 cell/tissue types were greater than those of positive silencers compiled in them (Figure 1E), suggesting that the positive silencers used for training and testing consist of only a small portion of active silencers in these cell/tissue types.

Although we have successfully used four epigenetic marks (CA, H3K4me1, H3K4me3 and H3K27ac) as features to predict functional states of our CRMs as enhancers[50], in this study we only used three of them (CA, H3K4me1 and H3K27ac) in our LR model. We excluded H3K4me3, since it is more likely associated with promoters than to enhancers[60, 61]. We pooled positive and negative sets compiled in each of the 67 human cell/tissue types used in our previous study[50] (Materials and Methods), resulting in a positive set containing a total 1,415,796 positive enhancers and a negative set with the same number of negative control sequences. We trained and evaluated the seven LR models using all the seven possible combinations of the three epigenetic marks as features by 10-fold cross validation. Of the three models using only one mark, model 4 using CA had the highest median AUROC 0.913, followed by model 1 using H3K4me1 (median AUROC=0.897) and model 2 using H3K27ac (median AUROC=0.866). Of the three models using two marks, model 3 using H3K4me1 and H3K27ac (H3K4me1&H3K27ac) obtained the highest median AUROC of 0.971, followed by model 5 (CA&H3K4me1, median AUROC=0.963) and model 6 (CA&H3K27ac, median AUROC=0.952). Model 7 using all of the three marks achieved the highest median AUROC of 0.977 (Supplementary Figure S1A and S1B), which is significantly higher than the other six models (p value < 0.01, Mann-Whitney U test). Consistently, CA has a higher weight (92.0) in the model than H3K4me1 (30.0) and H3K27ac (17.1) (Supplementary Figure S1C). The median AUROC value achieved by model 7 (0.977) is comparable with our previous model (0.986) using four epigenetic marks, which substantially outperforms five existing state-of-the-art methods[50]. We thus selected model 7 as our enhancer functional state predictor (enhancer predictor) for the subsequent predictions. As expected, the numbers of predicted active enhancers in 65 of the 67 cell/tissue types are greater than those of positive enhancers compiled in them (Supplementary Figure S1D), suggesting that the positive enhancers used for training and testing consist of only a small portion of active enhancers in most of the cell/tissue types. However, both the positive sets and predicted active enhancers in each of these cell/tissue type are smaller than those compiled and predicted in our earlier study[50], due to the more stringent criterion used to compile the positive sets to ensure the CRMs in the positive sets are true active enhancers.

In summary, CA alone is a more effective predictor for both active silencers (Figure 1B) and active enhancers (Supplementary Figure S1A) than the other two marks (H3K27me3 and H3K9me3 for silencers and H3K4me1 and H3K27ac for enhancers) alone. Using additional two histone marks could moderately improve the enhancer prediction accuracy (mean AUROC 0.977 vs 0.913), but only slightly increase the silencer prediction accuracy (mean AUROC 0.962 vs 0.948), over that obtained by using CA alone. In both predictors CA has overwhelmingly higher weights than the other two marks, making the other two marks weaker predictors.

**Figure 1.**
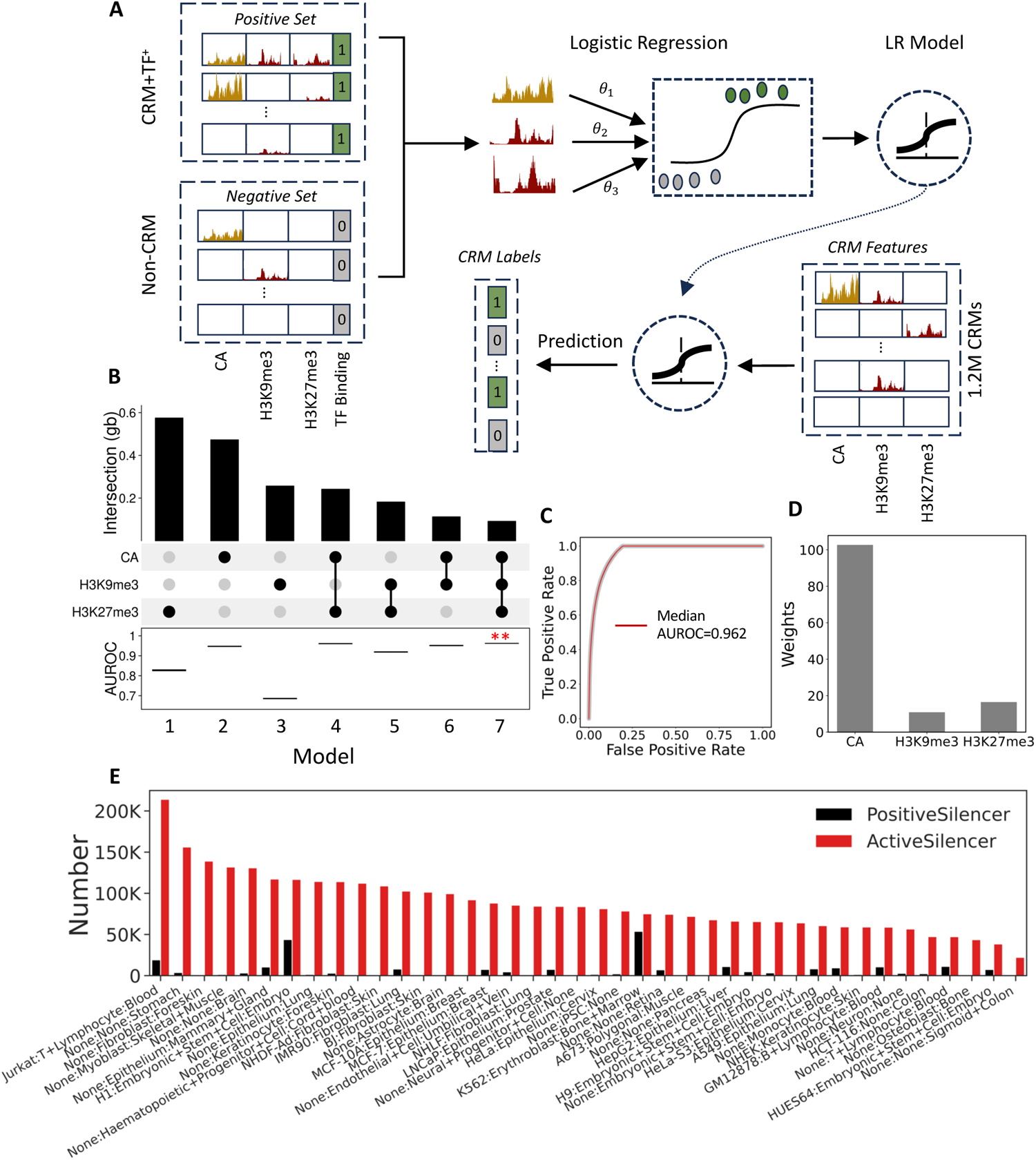
The epigenetic marks can accurately predict the functional states of putative silencers. **A.** A cartoon illustrating the workflow of our LR model using CA, H3K9me3 and H3K27me3 signals as the features. **B.** The UpSet plot showing intersection sizes (Gb) of mark peaks (upper bar graph) and the boxplot of AUROCs of the seven LR models using all possible combinations of the three epigenetic marks with 10-fold cross validation (lower boxplots). **: p value < 0.01, Mann-Whitney U test. **C.** ROC curves of model 7 using all the three marks. The red curve is the median ROC curve from the results of 10-fold cross validation. The AUROC curve of each fold is invisible since these curves have almost the same shape as the median curve. **D.** Bar graph of the weights of CA, H3K9me3 and H3K27me3 in model 7. **E.** Bar graph of the numbers of positive and active silencers compiled and predicted, respectively, in each of the 40 cell/tissue types.

### Varying portions of the 1.2M CRMs are active as enhancers or silencers in various cell/tissue types

Using our enhancer and silencer predictors trained on the pooled positive and negative sets in the 67 and 40 (Supplementary Table S1-S3) cell/tissue types, respectively, with the required epigenetic data available from Cistrome [62, 63], we predicted the functional states of our 1.2 M predicted CRMs[42] as enhancers in 105 cell/tissue types and as silencers in 58 cell/tissue types, respectively, with the required epigenetic data available from ENCODE [64] (Figure 2A). This yielded highly varying numbers of active enhancers in the 105 cell/tissue types ranging from 31,947 (2.7% of the 1.2 M CRMs) in tibial nerve cells to 168,471 (14.3%) in motor neuron cells, with a median of 95,496 (8.1%) in a cell/tissue type (Supplementary Table S4). We predicted a total of 10,068,782 active enhancers in the 105 cell/tissue types. After removing the redundancy, we ended up with a total of 695,507 (59.0%) non-redundant active enhancers from the 105 cell types. Moreover, we also predicted highly varying numbers of active silencers in the 58 cell/tissue types ranging from 27,843 (2.4%) in tibial nerve cells to 197,133 (16.7%) in HepG2 cells, with a median of 86,668 (7.4%) in a cell/tissue (Supplementary Table S5). We predicted a total of 5,096,269 active silencers in the 58 cell/tissue types. After removing the redundancy, we ended up with a total of 677,840 (57.5%) non-redundant active silencers from the 58 cell types. Thus, we predicted a slightly higher median number of active enhancers than active silencers (95,496 vs 86,668). As expected, most (78.0%∼97.2%) of the CRMs were not active either as enhancers or as silencers in a cell/tissue type (Supplementary Table S6). In total, we predicted the functional types and states for 868,944 (73.8%) of the 1.2 M CRMs as active enhancers or active silencers in at least one of the cell/tissue types.

### Predicted functional types and states of CRMs are reflected by their epigenetic mark signals

Notably, in each of the 49 cell/tissue types with only enhancer marks (CA, H3K4me1 and H3K27ac) data available (Materials and Methods), we were only able to predict each CRM either as an active enhancer or as an inactive enhancer (Figure 2A, Supplementary Table S7). For example, in the A549 cells, we predicted 128,601 (10.9%) CRMs to be active enhancers and the remaining 89.1% to be inactive enhancers (Supplementary Table S7). The predictions in each of these 49 cell/tissue types are reflected by the signal patterns of all the three epigenetic marks on the CRMs. Figure 2B shows the case in the A549 cells from donor ENCDO000AAZ as an example. Specifically, CRMs that were predicted to be active enhancers in a cell/tissue type such as the A549 cells were enriched in the active enhancer marks (CA, H3K4me1 and H3K27ac), while those that were not, were depleted of these signals (Figure 2B). Similarly, in each of the two cell/tissue types with only putative active silencer marks (CA, H3K27me3 and H3K9me3) data available (Materials and Methods), we were only able to predict each CRM either as an active silencer or as an inactive silencer (Figure 2A, Supplementary Table S8). For example, in the heart left ventricle cells from donor ENCDO039RUH, we predicted 108,302 (9.2%) CRMs to be active silencers and the remaining 90.8% to be inactive silencers (Supplementary Table S8). The predictions in these two cell/tissue types also are reflected by the signal patterns of all the three epigenetic marks on the CRMs. Figure 2C shows the case for the heart left ventricle cells. Specifically, CRMs that were predicted to function as active silencers in a cell/tissue type such as heart left ventricle cells were enriched in putative active silencer marks (CA, H3K27me3 and H3K9me3), while those that were not, were depleted of the signals (Figure 2C).

**Figure 2.**
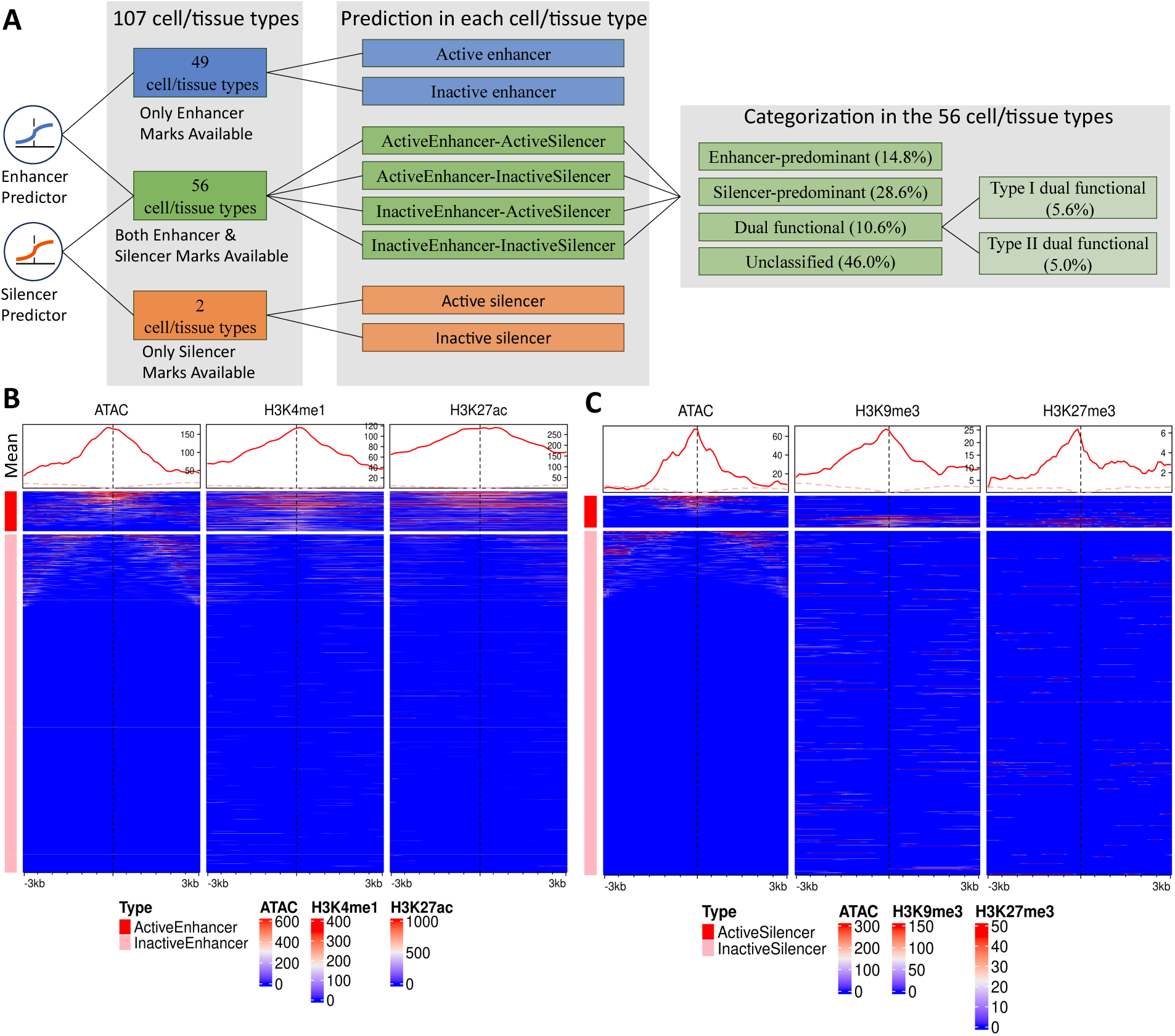
Prediction of functional types and states of CRMs in the 107 cell/tissue types. **A.** Prediction of functional types and states of CRMs in various cell/tissue types with different available data and categorization of CRMs based on predictions in the 56 cell/tissue types with both active enhancer and silencer marks data available. **B.** Heatmaps of signals of three active enhancer marks in a 6 kb window centering on the middle points of the predicted active enhancers and inactive enhancers in A549 cells from donor ENCDO000AAZ. **C.** Heatmaps of signals of three putative active silencer marks in a 6 kb window centering on the middle points of the predicted active silencers and inactive silencers in the heart left ventricle cells from donor ENCDO039RUH. The heatmaps show the mean signals of the epigenetic marks in each 100 bp sliding window along each sequence; the line plot shows the mean signal of each window at a position in all the sequences in the set (Materials and Methods). The color code for the types in the line plot above each column is the same as the left legends of the heatmaps.

Furthermore, in each of the 56 cell/tissue types with both enhancer and putative silencer marks data available (Materials and Methods), we have four possible predictions about the functional types and states of each of the 1.2M CRM (Figure 2A): i) both the enhancer predictor and the silencer predictor predict it to be active (ActiveEnhancer-ActiveSilencer); ii) the enhancer predictor predicts it to be active, but the silencer predictor predicts it to be inactive (ActiveEnhancer-InactiveSilencer); iii) the enhancer predictor predicts it to be inactive, but the silencer predictor predicts it to be active (InactiveEnhancer-ActiveSilencer); and iv) both the enhancer predictor and the silencer predictor predict it to be inactive (InactiveEnhancer-InactiveSilencer). The numbers of predicted CRMs in each of the categories are shown in Figure 3A (Supplementary Table S9). For example, in the MCF-7 cells from donor ENCDO000AAE, we predicted 69,327 (5.9%) CRMs to be “ActiveEnhancer-ActiveSilencer”, 45,278 (3.8%) to be “ActiveEnhancer-InactiveSilencer”, 45,862 (3.9%) to be “InactiveEnhancer-ActiveSilencer”, and the remaining 1.02M (86.4%) to be “InactiveEnhancer-InactiveSilencer” (Figure 3A, Supplementary Table S9). The predictions in each cell/tissue type also are reflected by the relevant epigenetic marks on the CRMs. Figures 3B and 3C show the cases in the MCF-7 cells as examples. Specifically, “ActiveEnhancer-ActiveSilencer” CRMs were enriched in both the marks of active enhancers (CA, H3K4me1 and H3K27ac) and marks of putative active silencers (CA, H3K9me3 and H3K27me3) (Figure 3B). “ActiveEnhancer-InactiveSilencer” CRMs were enriched in marks of active enhancers H3K4me1 and H3K27ac but depleted of marks of putative active silencer H3K9me3 and H3K27me3 (Figure 3B). Interestingly, the CA signals on these “ActiveEnhancer-InactiveSilencer” CRMs were weak in the middle but strong at the two flanking regions, while the H3K4me1 signals were narrowly peaked at the middle, suggesting that the middle of these CRMs might not be nucleosome free, and therefore could not be cut by transposase. “InactiveEnhancer-ActiveSilencer” CRMs were enriched in putative active silencer marks but depleted of active enhancer marks H3K4me1 and H3K27ac (Figure 3C). “InactiveEnhancer-InactiveSilencer” CRMs had weak signals of all the five epigenetic marks (Figure 3C). In summary, by predicting the functional states of the 1.2 M CRMs as enhancers or silencers in a cell/tissue type, we are able to simultaneously predict the functional states and types of the CRMs using only five epigenetic marks data in the very cell/tissue type.

**Figure 3.**
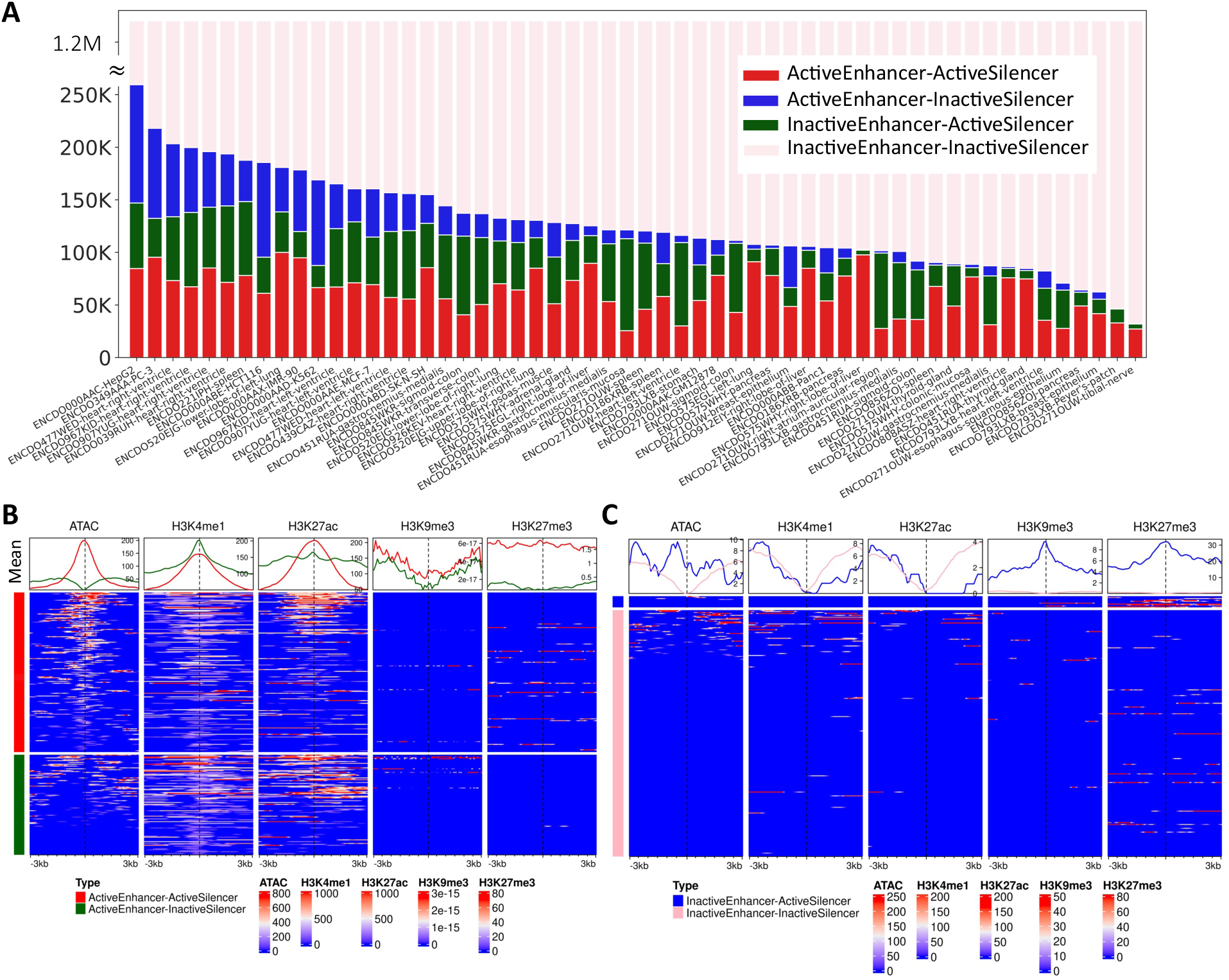
Four possible combinations of predictions of the functional types and states of the CRMs in the 56 cell/tissue types with both active enhancer and putative active silencer marks data available. **A.** Bar plots of the numbers of the CRMs with the four possible combinations of predicted functional types and states in each of the 56 cell/tissue types. **B.** Heatmaps of signals of the five epigenetic marks in a 6 kb window centering on the middle of the predicted ActiveEnhancer-ActiveSilencer and ActiveEnhancer-InactiveSilencer CRMs in MCF-7 cells. **C.** Heatmaps of signals of the five marks in a 6 kb window centering on the middle of the predicted InactiveEnhancer-ActiveSilencer and InactiveEnhancer-InactiveSilencer CRMs in MCF-7 cells. The heatmaps show the mean signals of the epigenetic marks in each 100 bp sliding window of each sequence; the line plot shows the mean signal of each window at a position of all the sequences in the set (Materials and Methods). The color code for the types in the line plot above each column is the same for the heatmaps.

### At least 10% of the CRMs are dual functional

We predicted a total of 793,140 (67.3%) CRMs to be active as enhancers and/or silencers in at least one of the 56 cell/tissue types (Figure 2A, Supplementary Table S9) with both active enhancer and putative silencer marks data available. These predictions provide us an opportunity to investigate the predominant roles of the CRMs used as enhancers, silencers, or both in these cell/tissue types. Of these 793,140 CRMs, 117,646 (14.8%) were predicted to be active only as enhancers across all the cell/tissue types (Enhancer-predominant), 227,211 (28.6%) were predicted to be active only as silencers across the cell/tissue types (Silencer-predominant), and 448,283 (56.6%) were predicted to be active both as enhancers and silencers in the 56 cell/tissue types (Dual functional CRMs) (Figure 2A). Of the 448,283 dual functional CRMs, 408,451 (91.1%) were predicted to be both as active enhancers and active silencers in the same cell/tissue types (denoted as type I dual functional CRMs), while the remaining 39,832 (8.9%) were predicted to be as active enhancers in some cell/tissue types and as active silencers in other cell/tissue types (denoted as type II functional CRMs). As we indicated earlier, CA signals have much higher weights than the two other markers in both our enhancer (Supplementary Figure 1C) and silencer (Figure 1D) predictors, thus a CRM with a very strong CA signal but relatively weak signals of the two other marks can be predicted both as active enhancer and as active silencer in the same cell/tissue type. To reduce possible false positives, for type I dual functional CRMs, we only consider those that have at least one active enhancer mark and at least one putative active silencer mark for further analysis, yielding a total of 44,153 (10.8%) more stringent type I dual functional CRMs, while the remaining 89.2% were categorized as unclassified CRMs (Figure 2A). Our subsequent analyses will mainly focus on a total of 83,985 (10.6%) dual functional CRMs (44,153 type I and 39,832 type II) (Figure 2A).

### Dual functional CRMs can switch their roles in different cellular contexts

It has been shown in previous reports that CRMs may switch their roles between active enhancers and active silencers in different cellular contexts[9, 10, 56]. Consistently, our type II dual functional CRMs functioned as active enhancers in some cell/tissue types while as active silencers in other cell/tissue types. On the other hand, all of the 44,153 type I dual functional CRMs could function both as active enhancers and as active silencers in at least one of the 56 cell/tissue types, most (89.1%) of which could also only function as active enhancers or active silencers in at least one of the 56 cell/tissue types. Moreover, more than half (56%) of the type I dual functional CRMs have dual functions in the same cell/tissue type in only one cell/tissue type, and only a small portion (3.8%) of them did so in more than half (28) of the 56 cell/tissue types (Figure 4A). Thus, most type I dual functional CRMs also were able to switch their roles in different cell/tissue types. For example, the CRM at chr7:1,428,212-1,430,677 was dual functional in spleen (ENCDO221IPH) and HepG2 (ENCDO000AAC) cells, but only functioned as an active enhancer in Panc1 (ENCDO000ABB) cells, and only functioned as an active silencer in heart left ventricle (ENCDO477WED), SK-N-SH (ENCDO000ABD) and breast epithelium (ENCDO271OUW) cells, as indicated by the patterns of epigenetic marks on the CRM in relevant cell/tissue types. More specifically, as shown in Figure 4B as examples, in spleen cells the CRM was heavily marked by both active enhancer marks (H3K4me1) and active silencer marks (H3K27me3) in addition to CA, while in Panc1 cells it was heavily marked by the active enhancer marks (H3K4me1 and H3K27ac), but depleted of active silencer marks, and in heart left ventricle cells it was heavily marked by the active silencer marks (H3K27me3), but with weak active enhancer marks.

**Figure 4.**
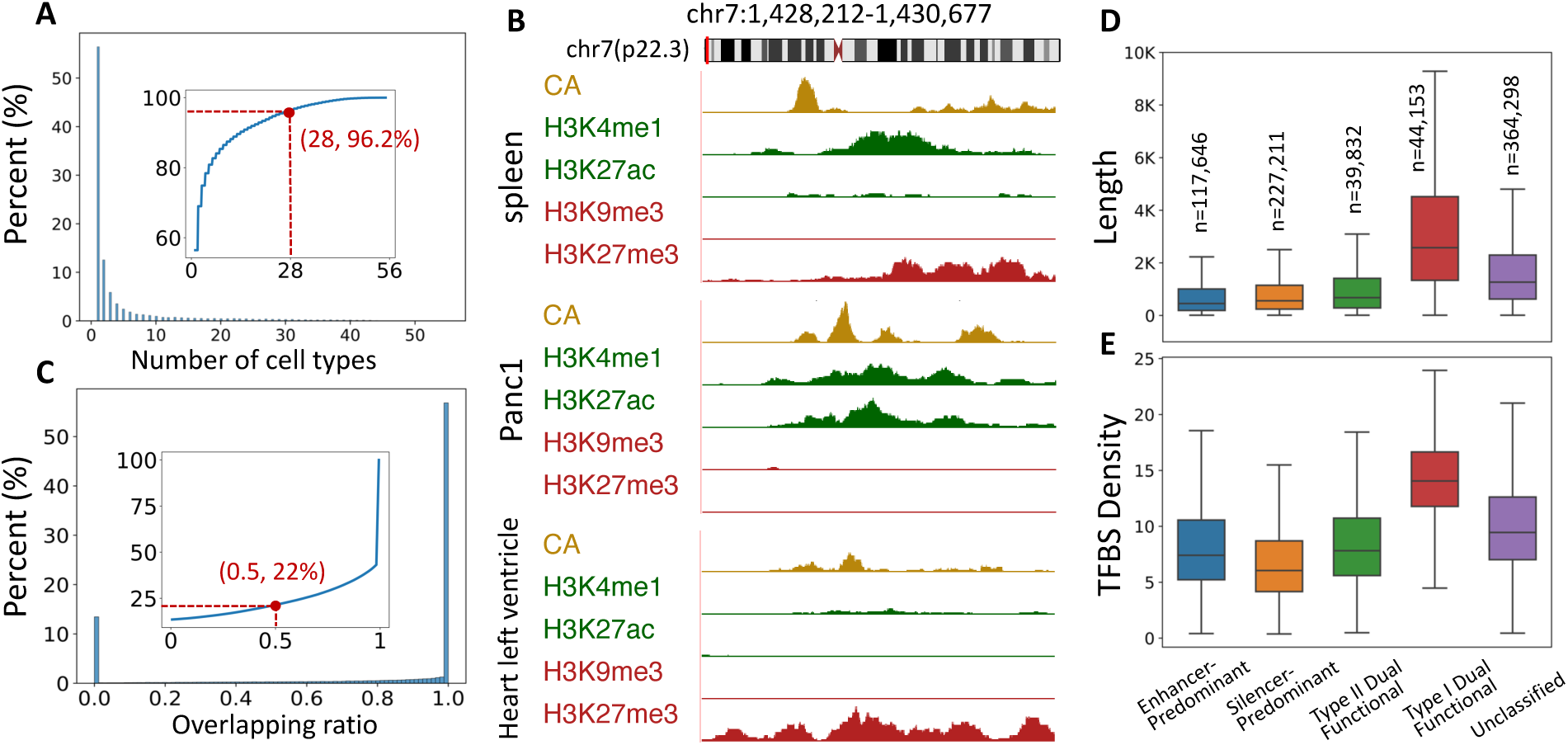
Analysis of different types of active CRMs in the 56 cell/tissue types. **A.** Histogram of percentages of type I dual functional CRMs that are active both as enhancers and silencers in the indicated number of cell/tissue types. The inset plot is cumulative percentage of type I dual functional CRM that are active both as enhancers and silencers in at least the indicated numbers of cell/tissue types. **B.** Epigenetic marks on a type I dual functional CRM at chr7:1,428,212-1,430,677 in spleen cells where it functions both as an active enhancer and as an active silencer (upper panel); in Panc1 where it functions only as an active enhancer (middle panel); and in heart left ventricle cells where it functions only as an active silencer (lower panel). **C.** Histogram of percentages of type I dual functional CRMs with the indicated overlapping ratio between their enhancer mark regions and silencer mark regions. The inset plot is the cumulative percentage of overlapping ratio less than the indicated numbers. **D.** Boxplots of the lengths of the four types of CRMs and unclassified CRMs. **E.** Boxplots of the density of TFBSs (number of TFBSs per 100 bp) in the four types of CRMs and unclassified CRMs.

### Enhancer and silencer mark peaks on type I dual functional CRMs overlap each other

As we predicted CRMs by stitching adjacent TFBSs[42], it is conceivable that type I dual-functional CRMs may be the result of simply merging distinct enhancers and silencers. To test this possibility, we assessed the extent to which active enhancer mark peaks and putative active silencer mark peaks along type I dual functional CRMs overlap each other (Materials and Methods). As shown in Figure 4C, of type I dual functional CRMs, 55% have their active enhancer and silencer mark peaks completely overlapping each other (overlapping ratio = 1), and more than 78% have an overlapping ratio larger than 0.5, indicating that type I dual functional CRMs are unlikely formed by incorrectly stitching adjacent enhancers and silencers. For example, active enhancer and silencer marks interdigitate and overlap one another along the CRM chr7:1,428,212-1,430,677 in the spleen cells where it functions both as an active enhancer and as an active silencer (Figure 4B). These results indicate that the dual functions of a CRM might be achieved by the collaboration between different parts of the CRM, but not by two non-overlapping parts each conferring the CRM a different role. It is likely that such collaboration renders the dual functional CRMs to be longer than enhancer-predominant and silencer-predominant CRMs.

### Length and TFBS density of a CRM reflect the complexity of its functional type

To see how the length of a CRM is related to its predicted functional type, we plotted the distributions of the lengths of the different types of predicted CRMs. Interestingly, as shown in Figure 4D, different types of CRMs show distinct length distributions. Specifically, type I dual functional CRMs have the longest median length (2,577 bp), followed by type II dual functional CRMs (678 bp), silencer-predominant CRMs (558 bp), and enhancer-predominant CRMs (454 bp). Unclassified CRMs are shorter than type I dual functional CRMs, yet longer than the other three types, suggesting that they might be a blend of type I dual functional CRMs and other three types of CRMs. The longer lengths of dual functional CRMs might be related to their more complex functions. Moreover, it has been demonstrated that enhancers and silencers in mouse retinal photoreceptor cells (cones and rods) possess different information content in terms of TFBS composition[65]. In light of this, we analyzed TFBS densities (number of TFBSs in 100 pb length, Materials and Methods) of enhancer-predominant CRMs, silencer-predominant CRMs, type I and type II dual functional CRMs, and unclassified CRMs. As shown in Figure 4E, type I dual functional CRMs have the highest TFBS densities, followed by type II dual functional CRMs, enhancer-predominant CRMs and silencer-predominant CRMs. Likewise, unclassified CRMs exhibit a lower TFBS density than type I dual functional CRMs, but a higher TFBS density than the other three types (Figure 4E), suggesting again that unclassified CRMs might be a combination of type I dual functional CRMs and other three types of CRMs.

### Type I dual functional CRMs might execute dual functions by regulating different genes in the same cell type

Of the 56 cell/tissue types that we used to predict type I dual functional CRMs (Figure 2A), 9 are cell lines, each nominally contains a single cell type; while the remaining 47 are primary tissues, each might contain multiple cell types. Thus, we compared the numbers of type I dual functional CRMs that functioned as both active enhancers and active silencers in the cell lines (n=9) with those in the primary tissues (n=47). As shown in Figure 5A, the numbers of dual active CRMs observed in the cell lines were not significantly different (p-value>0.26) from those in the primary tissues. This suggests that the dual functions of CRMs may not necessarily be attributed to various cell types in a primary tissue, rather, CRMs can be dual functional in the same cell type.

To investigate how dual functional CRMs could possibly exert their enhancer and silencer functions, we identified genes whose promoters were in close physical proximity to a dual functional CRMs from the Hi-C interaction map and compared expression levels of putative target genes in different cell/tissue types according to the CRM’s predicted functional states as an enhancer or a silencer of the genes. For instance, the Hi-C interaction map shows that CRM chr7:1,428,212-1,430,677 is in close physical proximity with the promoters of genes *MICALL2, INST1, MAFK,* and *PSMG3* (Figure 5B). Figure 5C shows the expression levels of these genes across diverse cell/tissue types based on the CRM’s functional states as an enhancer for these genes. *INST1* and *PSMG3* had significantly higher expression levels in cell/tissue types where the CRM was active as an enhancer than in cell/tissue types where the CRM was inactive as an enhancer, while *MICALL2* and *MAFK* did not. This leads us to conclude that the CRM might function as an enhancer for *INST1* and *PSMG3*, but not for *MICALL2* and *MAFK*. Similarly, Figure 5D shows the expression levels of the four genes across different cell/tissue types based on the CRM’s functional states as a silencer for the genes. *MICALL2* had significantly lower expression levels in cell/tissue types where the CRM was active as a silencer than in cell/tissue types where the CRM was inactive as a silencer, while the other three genes did not. We therefore conclude that the CRM might function as a silencer for *MICALL2*, but not for the other three genes. Although the CRM interacts with gene *MAFK*, our result suggests that it may not regulate this gene as either an enhancer or a silencer in the cell/tissue types that we examined.

**Figure 5.**
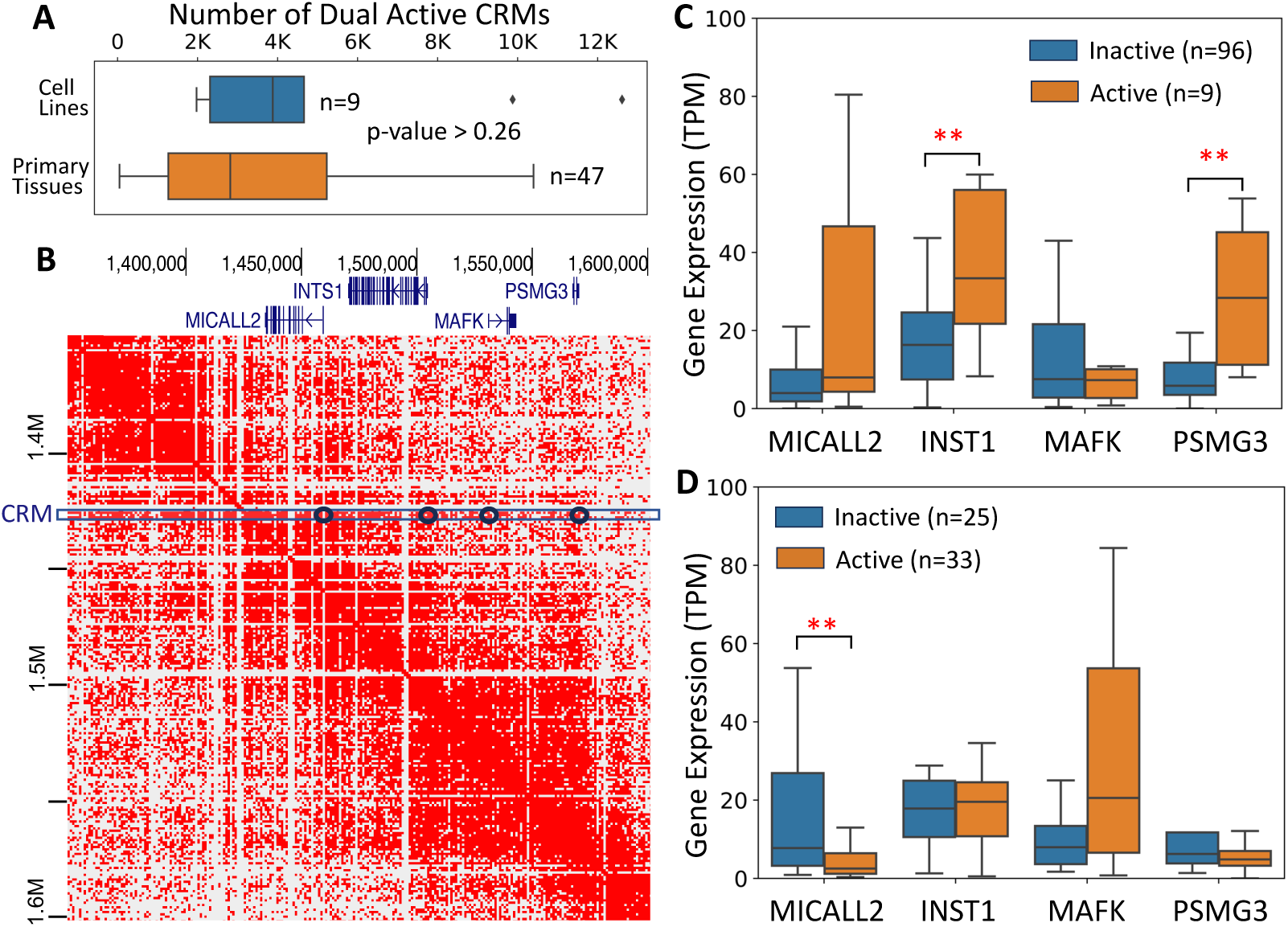
Comparison of numbers of type I dual functional CRMs in cell lines and primary tissues and an analysis of expression levels of putative target genes of a type I dual functional CRM. **A.** Boxplots of numbers of type I dual functional CRMs predicted in 9 cell lines and 47 primary tissues. p-value>0.26 (Mann-Whitney U-test). **B. A** Hi-C interaction map for the region of 1,350,000 to 1,60,000 bp on chromosome 7. CRM chr7:1428212-1430677 indicated by the box interacts with the promoters of genes *MICALL2, INST1, MAFK* and *PSMG3*, highlighted by the circles. **C.** Boxplots of expression levels of genes *MICALL2, INST1, MAFK* and *PSMG3* across different cell/tissue types based on the CRM’s functional states as an enhancer in these cell/tissue types. **: p-value <0.01 (Mann-Whitney U-test). **D.** Boxplots of expression levels of genes *MICALL2, INST1, MAFK* and *PSMG3* across different cell/tissue types based on the CRM’s functional states as a silencer in these cell/tissue types. **: p-value <0.01 (Mann-Whitney U-test).

### The *“validated” silencers* from silencerDB may contain false positives

We have previously shown that our CRM functional state predictor that used four active enhancer marks substantially outperformed five state-of-the-art methods[50]. As our enhancer predictor trained on a more stringent positive set using three active enhancer marks achieved comparable AUROC value (0.980) to that of our previous predictor (0.986) in the same dataset, to avoid repetition, here we only evaluated the performance of our silencer predictor. To this end, we first compared our 677,840 (57.5%) predicted silencers pooled from the 58 cell/tissue types (Figure 2A) with the “validated” silencers from the silencerDB database[66]. There were 8,588 “validated” unique silencers in silencerDB, which were identified by two recent studies using MPRA[58] or its variant called repressive ability of silencer elements (ReSE) screen[59]. Our predicted silencers overlap 4,661 (54.3%) of the “validated” silencers by at least a 1bp, while only 5,525 (64.3%) of the “validated” silencers overlap 5,434 (0.5%) of our predicted 1.2 M CRMs. To see whether the rest 3,063 (35.7%) that do not overlap our CRMs are really functional, we analyzed their evolutionary behaviors using the phyloP scores. As shown in Figure 6A, like our predicted silencers the “validated” silencers that at least partially overlap our CRMs are under strong selection as indicated by their broadly distributed phyloP scores [67]. By contrast, the remaining “validated” silencers that do not overlap our CRMs are largely selectively neutral or nearly so as indicated by their narrowly distributed phyloP scores around 0 (Figure 6A) [67]. We thus posit that the “validated” silencers that do not overlap our CRMs might represent false positives. If we exclude these false negative “validated” silencers (35.7%) and only consider the 5,525 (64.3%) of the validated silencers that overlap our predicted 1.2 M CRMs, then we recall 84.4% (4,661) of them. Thus, our method has achieved 84.4% sensitivity for recalling validated silencers overlapping our 1.2 M CRMs, substantially higher than by chance (0.3%=57.5% x 0.5%).

### Comparison of our predicted silencers with *predicted silencers* from two existing methods

Next, we compared our 677,840 silencers with the predicted silencers from silencerDB, primarily by two previous studies[57, 59]. Specifically, Huang et al.[57] predicted a set of silencers by correlating putative active silencer epigenetic marks (H3K27me3) signals on DHSs with mRNA levels of neighboring genes across 25 different cell/tissue types, and then predicted additional silencers using an SVM model trained on the set using a combination of sequence features, epigenetic marks and gene expression profiles (CoSVM). Hawkins et al.[59] predicted silencers using a gkmSVM model by employing a simple subtractive strategy to obtain uncharacterized regulatory elements as potential silencers. After removing the redundancy in different cell/tissue types, we ended up with 157,813 and 982,985 non-redundant putative silencers predicted by CoSVM and gkmSVM, respectively.

**Figure 6.**
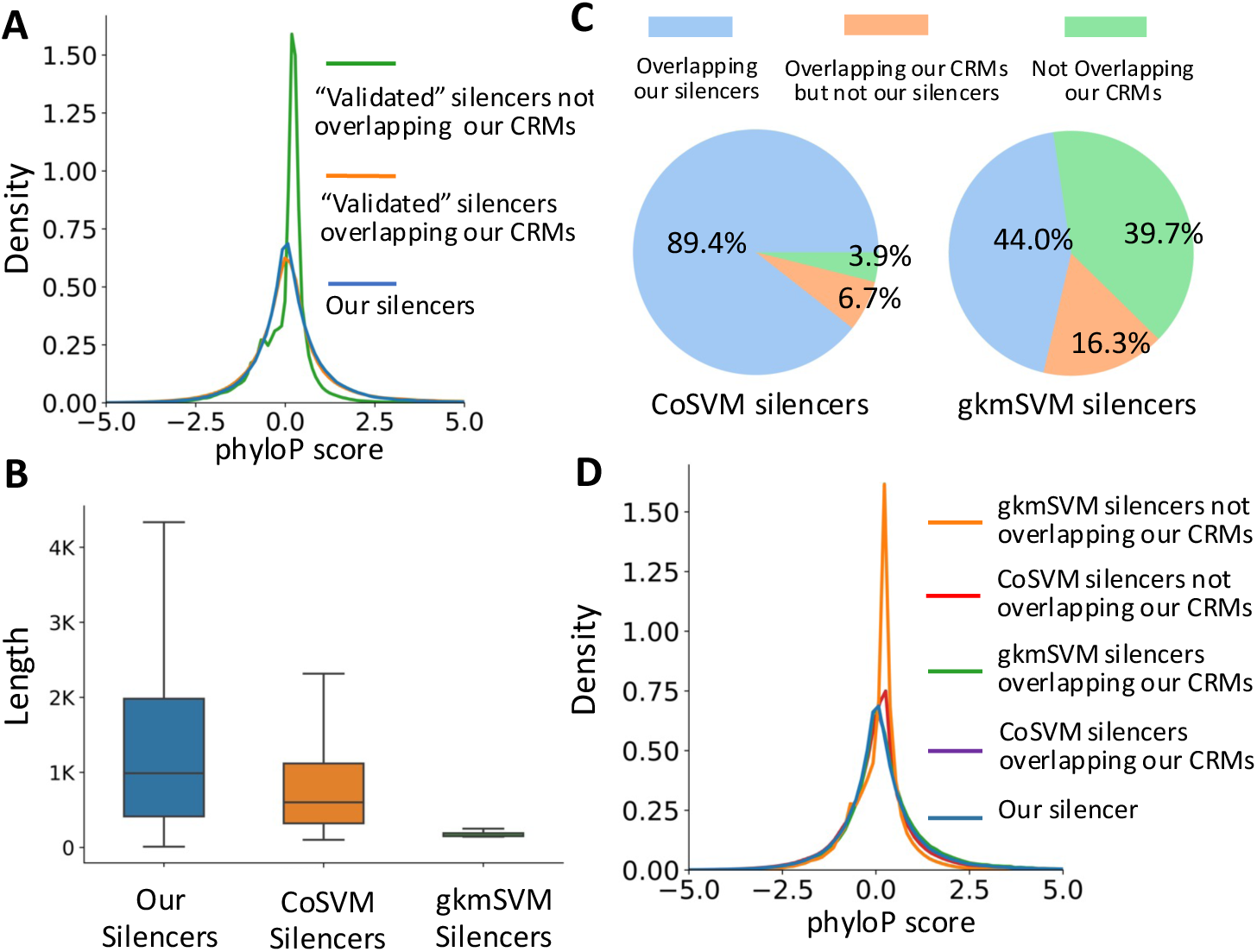
Comparison of our predicted silencers with the “validated” silencers and predicted silencers by CoSVM and gkmSVM. **A.** Distributions of phyloP scores of “validated” silencers that overlap or do not overlap our CRMs, as well as of our silencers. **B.** Boxplots of lengths of our predicted silencers, CoSVM-predicted silencers and gkmSVM-predicted silencers. **C.** Pie charts of CoSVM-predicted silencers (left) and gkmSVM-predicted silencers (right), which overlap our silencers, do not overlap our silencer but overlap our CRMs and do not overlap our CRMs. **D.** Distributions of phyloP scores of gkmSVM-predicted silencers and CoSVM-predicted silencers, which overlap and do not overlap our CRMs, as well as of our silencers.

As summarized in Table 1, gkmSVM predicted the highest number (982,985) of silencers, followed by our method (677,840) and CoSVM (157,813). However, our predicted silencers with a median length of 989 bp are longer than those predicted by CoSVM (601 bp) and gkmSVM (150 bp) (Figure 6B) and covers a greater proportion (33.3%) of the genome than those by CoSVM (4.8%) and gkmSVM (5.7%) (Table 1). Only 38,525 (24.4%) of the CoSVM-predicted silencers could be mapped to 52,539 (5.3%) gkmSVM-predicted silencers, indicating that the two methods predicted quite different sets of sequences as silencers. On the other hand, of the CoSVM-predicted silencers, 151,706 (96.1%) overlap 142,136 (12.1%) of our predicted 1.2 M CRMs, of which 141,062 (89.4%) overlap 127,380 (10.8%) of our predicted silencers, while 10,644 (6.7%) overlap our CRMs but do not overlap our silencers, and 6,107 (3.9%) do not overlap our predicted CRMs (Figure 6C). As expected, the 151,706 CoSVM-predicted silencers that overlap our CRMs have similar evolutionary constraints as our predicted silencers (Figure 6D), suggesting that they might be true silencers. In this regard, our predicted silencer recalled most (93.0%) of CoSVM-predicted silencers, substantially higher by chance (7%=57.5% x 12.1%). Interestingly, the 6,107 CoSVM-predicted silencer that do not overlap our CRMs also have similar evolutionary constraints as our predicted silencers (Figure 6D), suggesting that they might be true silencers, but our CRM predictor dePCRM2 failed to predict them to be CRMs. This could be explained by the fact that dePCRM2 is not able to touch about 15% of the genome because of unavailability of TF binding data in these regions, therefore missing those CRMs that can function as silencers.

Moreover, of the gkmSVM-predicted silencers, 592,502 (60.3%) overlap 342,813 (29.1%) of our 1.2 M predicted CRMs, 432,572 (44.0%) overlap 232,960 (19.8%) of our predicted silencers, 159,930 (16.3%) overlap our CRMs but do not overlap our silencers, and 390,483 (39.7%) do not overlap our predicted CRMs (Figure 6C). As expected, the 592,502 gkmSVM-predicted silencers that overlap our CRMs have similar evolutionary constraints as our predicted silencers (Figure 6D), suggesting that they might be true silencers. In this regard, our method recalled most (73.0%) of them, substantially higher by chance (16.7%=57.5% x 29.1%). In contrast, the gkmSVM-predicted silencers that do not overlap our CRMs are largely selective neutral or nearly so, suggesting that they are more likely false positives (Figure 6D). In summary, CoSVM-predicted silencers appear highly accurate, and our predicted silencers recall 93.0% of them. Of the gkmSVM-predicted silencers, 44.0% appear to be authentic, and our predicted silencers recall 73.0% of them, while 39.7% appear to be false positives. Our method is comparable to CoSVM but superior to gkmSVM in accuracy. However, our method predicts more silencers than CoSVM.

**Table. 1.** Summary of silencers predicted by the three methods.

| Method | # of unique silencers | Genome coverage | Median length |
| --- | --- | --- | --- |
| CoSVM[57] | 157,813 | 4.75% | 601 bp |
| gkmSVM(ENCODE)[59] | 982,985 | 5.67% | 150 bp |
| Our prediction | 677,840 | 33.3% | 989 bp |

## Discussion

In this study, we introduce two LR models to separately predict the functional states of our previously predicted 1.2M CRMs[42] as enhancers and silencers. The enhancer predictor uses signals of three epigenetic marks (CA, H3K4me1 and H3K27ac) on the CRMs as features. We choose these marks since they have been shown to be associated with active enhancers[68]. The silencer predictor also employs signals of three epigenetic marks (CA, H3K9me3 and H3K27me3) on the CRMs as features. We choose these three marks based on the following reasons: CA is a hallmark of TF binding on CRMs including silencers[69, 70], both H3K9me3[71, 72] and H3K27me3[73] have been shown to be associated with repressive DNA sequences, and CA as well as H3K27me3 have been used as features for predicting silencers[57]. As many cell/tissue types only have one set of these epigenetic marks, we build two independent predictors for their wider applicability as demonstrated in this study. Our enhancer predictor achieves comparable AUROC (0.977 vs 0.986) as our previous predictor that used four marks, which substantially outperforms five state-of-the-art methods[74]. Our silencer predictor also achieves a high AUROC of 0.962, albeit slightly smaller than that (0.977) of the enhancer predictor. Although none of the three epigenetic marks alone or their combinations are specific for either active enhancers[45, 46, 49] or silencers[57], each of the three marks alone and their combinations achieve from moderate (0.685) to high (>0.95) AUROC values. We attribute the high accuracy of the models to our two-step approach[74], i.e., we predict the functional types and states (second step) of our CRMs that were predicted (first step) using TF binding data. This conclusion is in good agreement with earlier reports that once the loci of CRMs are accurately anchored by the binding of key TFs, epigenetic marks can be accurate predictors of the functional states of the CRMs[45, 46, 49]. Applying the enhancer model to the 105 cell/tissue types with data of the three active enhancer marks available, and the silencer model to the 58 cell/tissue types with data of the three putative active silencer marks available, we predict 868,944 (73.8%) of our 1.2 M CRMs in the human genome[42] to be active as enhancers or silencers in at least one of the 107 cells/tissue types.

Particularly, in the 56 of the 107 cell/tissue types, with both active enhancer and silencer marks data available, we predict 793,140 (67.3%) CRMs to be active enhancers, active silencers, or both in at least one of the cell/tissue types. We classify the 793,140 CRMs in four types: enhancer-predominant, silencer-predominant, dual functional, and unclassified. Moreover, we further classified dual functional CRMs into type I and type II based on whether or not they can be both active enhancers and active silencers in the same cell/tissue type. Moreover, since CA has much higher weights than the two other marks in both the enhancer and silencer predictors, they may predict some CRMs with very high CA signals but low signals of the two other marks both as active enhancers and silencers in the same cell/tissue type, thereby overestimating dual functional CRMs. To reduce possible such false positive predictions, we only consider the CRMs that are predicted to be active by both predictors as type I dual functional CRMs only if they are also labeled by at least one of the two other enhancer marks and one of the two other silencer marks, and classify the remaining CRMs that do not meet this criterion as unclassified. This gives a lower bounder 10.6% of the 793,140 CRMs to be dual functional. Consistent with earlier reports[9, 54, 56], type II dual functional CRMs may switch their roles in different cell/tissue types, presumably by binding two different sets of TFs in different cellular contexts. To the best of our knowledge, we for the first time find that type I dual functional CRMs can function both as enhancers and silencers in the same cell/tissue type.

We show that the four types of CRMs (enhancer-predominant, silencer-predominant, type I dual functional and type II dual functional) possess distinct properties in terms of their lengths and TFBS densities, which reflect their functional complexity. Specifically, the longer lengths and the higher TFBS densities of type I dual functional CRMs might also be related to their dual functions in the same cell/tissue type, necessitating longer length and denser TF bindings. The shorter lengths and lower TFBS densities of type II dual functional CRMs than those of type I dual functional CRMs might be due to the fact that TFBSs in the former type can be shared for different functions across different cell/tissue types, since they only serve a single function in each cell/tissue type, while this might not be the case for the latter type. Such sharing might result in reduced TFBS densities and shorter lengths of type II dual functional CRMs. The higher TFBS density of type II dual functional CRMs than those of enhancer-predominant and silencer-predominant CRMs suggest that the former type might need denser TFBSs than the latter two types to execute both enhancer and silencer functions in different cell/tissue types by interacting with different sets of TFs. Although enhancer-predominant CRMs tend to be shorter than silencer-predominant CRMs (Figure 4D), the higher TFBS density of the former type suggests that activating genes might be a more intricate process than repressing genes. Unclassified CRMs with intermediate lengths and TFBS densities might be a mixture of the four types of CRMs. In the future, we need to determine the types of unclassified CRMs. One possible approach is to consider the positive or negative correlation between the predicted activation probabilities of a CRM and the expression levels of its potential genes in a TAD across multiple cell/tissue type, as well as the physical proximity between the CRM and the transcription start sites of the potential genes as we demonstrated for the CRM shown in Figure 5. Alternatively, we may use more specific epigenetic marks for active enhancers and silencers as features in machine-learning models when such marks are available in the future.

The substantial or complete overlaps between enhancer and silencer epigenetic marks peaks along type I dual functional CRMs strongly suggest that they are not artifacts by erroneously concatenating enhancers/silencers with their adjacent silencers/enhancers. A type I dual functional CRM might accomplish its dual roles in the same cell by simultaneously binding two sets of TFs via their cognate binding sites that are interdigitated along the CRM. Alternatively, it might accomplish its dual roles in different individual cells in a cell population of the same type by separately binding two sets of TFs in different individual cells. However, before relevant single-cell data are available, we could not differentiate these two possibilities. In either scenario, the high overlaps between the epigenetic mark peaks of enhancers and silencers along type I dual functional CRMs suggest that they might fulfill their dual roles by collaborative bindings of two different sets of TFs to their cognate binding sites along the CRMs as recently suggested for enhancers[75]. Although type I dual functional CRMs can function both as enhancers and silencers in the same cell types, they more often only function as enhancers or as silencers in different cell/tissue types, presumably by binding one of the two sets of TFs, indicating the highly dynamic and cellular context dependent nature of their usage.

Our predicted silencers recall 84.4% of MPRA-validated silencers falling in our predicted CRMs, while missing the remaining 15.6%, indicating that we might need data from more cell/tissue types to predict them. On the other hand, we find that the “validated” silencers that do not overlap our predicted CRMs might be false positives as they are largely selective neutral. This result is consistent with our recent finding[76] and other reports[77–81] that a considerable proportion of “regulatory sequences” identified by expression vector-based methods such as MPRA and its variants might be false positives. The two sets of previously predicted silencers cover a similar proportion (4.75% vs 5.67%) of the genome, but differ largely in their numbers (157,813 vs. 982,984) and have few overlaps. We find that CoSVM-predicted silencers are rather accurate, while gkmSVM-predicted silencers might have a false positive rate at least 39.7%. Our predicted silencers recall 93.0% CoSVM-predicted and 73.0% of gkmSVM-predicted silencers falling in our CRMs. Clearly, to recall the missed 7.0% of CoSVM-predicted and 27% of gkmSVM-predicted silencers falling in our CRMs, we might need more data from more diverse cell/tissue types. On the other hand, both CoSVM and gkmSVM miss 89.2% and 79.2% of our predicted silencers. Therefore, our method predicts much more silencers than CoSVM and gkmSVM while achieving accuracy comparable to that of CoSVM and superior to that of gkmSVM.

With the ongoing expansion of epigenetic and TF binding data available in a wide spectrum of cell/tissue types, our two-step approach holds the potential to uncover a more comprehensive map of CRMs in the genome, and then predict their functional types and states within these cell/tissue types. Based on the accurately predicted functional states of the CRMs and the expression levels of genes across a large number of cell/tissue types, as well as physical proximity between the CRMs and genes in TADs, it is possible to predict the target genes of the CRMs. This forward-looking perspective underscores the adaptive nature of our approach and its ability to yield deeper insights into the regulatory genomes and transcriptional machineries as datasets continue to grow and diversify in the future.

## Materials and Methods

### The datasets

We obtained a set of 1,178,229 predicted CRMs in the human genome from our prior work[42]. We downloaded histone mark ChIP-seq, DNase-seq, ATAC-seq and TF ChIP-seq data in 67 human cell/tissue types from Cistrome Datasets Browser[62, 63] (Supplementary Table S1-S3). All these 67 cell/tissue types have data for the three active enhancer marks (CA, H3K4me1 and H3K27ac), and 40 of them also have data for the three active silencer marks (CA, H3K27me3 and H3K9me3). We downloaded ATAC-seq and histone mark ChIP-seq data and RNA-seq data in 107 cell/tissue types from ENCODE data portal[64], of which 105 cell types have data for active enhancer marks (CA, H3K4me1 and H3K27ac), 58 have data for active silencer marks (CA, H3K27me3 and H3K9me3), 56 have data for both the active enhancer and active silencer marks, 49 only have data for active enhancers, and 2 only have data for active silencer marks (Figure 2A, Supplementary Table S10, S11). We downloaded the Hi-C contact matrix of the K562 cell line from the ENCODE[64] portal (https://www.encodeproject.org/files/ENCFF080DPJ/).

### Epigenetic mark feature scores

For each epigenetic mark *e* and a sequence *c*, which can be a CRM or non-CRM, we define a raw feature score as:

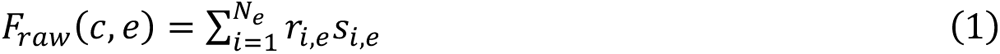

where *N_e_* is the number of peaks of *e* mapping to *c* at least 50% of the length of either one, *r_i,e_* the ratio of overlapping length between *c* and the *i_th_* peak of *e* over the length of the *i_th_* peak of *e*, *s_i,e_* the signal of the *i_th_* peak of *e* quantified by MACS2[82, 83]. We then normalized each raw epigenetic feature score in each cell/tissue type by the min-max normalization, i.e.,

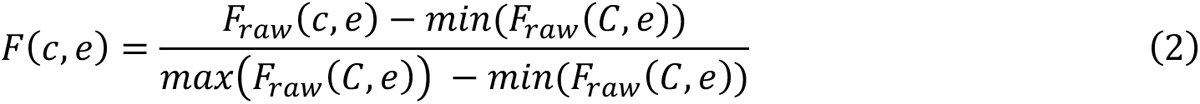

where *C* denotes all candidate sequences in the genome, 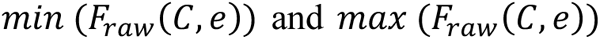 the minimum and maximum raw score of the epigenetic mark *e* over *C* in the cell/tissue type.

### Prediction of functional states of CMRs

Since a CRM can function both as an enhancer and a silencer in different cellular contexts, we use two separate models to predict the activation probability of a candidate CRM to be an enhancer and a silencer in a cell/tissue type. Specifically, we used an LR model to predict the activation probability of a candidate CRM functioning as an enhancer using signals of three active enhancer markers, i.e., CA, H3K4me1 and K3K27ac. Meanwhile, we used a similar LR model to predict the activation probability of a candidate CRM functioning as a silencer using signals of three active silencer markers, i.e., CA, H3K9me3 and K3K27me3.

#### Construction of positive and negative sets

In each of 67 cell/tissue types with the required data available, we selected the CRMs that overlap TF binding peaks and at least one of active enhancer marks H3K4m1 and K3K27ac, or of silencer marks K3K27me3 and K3K9me3 in the cell/tissue type as the positive enhancer or silencer set, to ensure the high quality of the positive set in a cell/tissue type. At the same time, we randomly selected predicted non-CRMs with matched numbers of the positive sets as the negative sets. We pooled the positive and negative sets in all the relevant cell/tissue types separately to construct a comprehensive positive set and a negative set for enhancers and silencers, resulting in 1,415,796 positive enhancers and 256,766 positive silencers and the same numbers of respective negative sets. Thus, the positive sets and negative sets for both enhancers and silencers are well-balanced.

#### Model training and evaluation

Ten-fold cross-validation was conducted to train and assess the performance of seven models using all the seven possible combinations of three marks as the features. The models were implemented using sci-kit learn v.0.24.2 and the code is available at https://github.com/zhengchangsulab/EnhancerSilencerPrediction.

#### Prediction

We applied both trained enhancer model and silencer model to the 1,178,229 CRMs in each of the 107 cell/tissue types with the required data available. We predict a CRM to be an active enhancer or silencer if its activation probability as an enhancer or a silencer is greater than 0.5.

### Heat maps of epigenetic marks

We used the “EnrichedHeatmap” package[84] version 4.2.2 to generate heat maps of signal intensities of epigenetic marks in a 6 kb region centered on a CRM. We computed the mean signal value for a mark in each100 bp sliding window in each of the 6 kb sequences, using “w0” as the “mean_mode”. The line plot on the top of the heat map is the mean signals of each window at a position across all the sequences of a set of CRMs. The CRMs within each set are ranked based on their CA signal strengths in descending order.

### Overlapping ratio of the enhancer and silencer epigenetic marks along dual functional CRMs

We merged the peak regions of the two enhancer marks H3K4me1 and H3K27ac on a CRM to form a unified region *E_mark_*, and those of the two silencer marks H3K9me3 and H3K27me3 on the CRM to form another unified region *Smark*. We computed the overlapping ratio between enhancer and silencer marks on the CRM as:

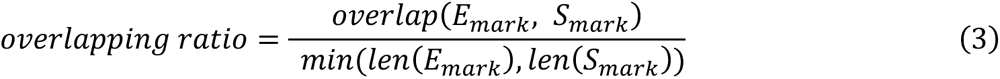

where *overlap*(*E_mark, S_mark*) denotes the length of the overlapping part between *E_mark_*_-_ and *S_mark_*, and *len(E_mark) and len(S_mark) t*he lengths of S*_mark_*, and *S_mark_*, respectively.

### Heat map of Hi-C contact matrix

We used the Juicebox[85] version 2.17.00 to generate the heat map of the Hi-C contact matrix using the Hi-C data from K562 cells with default settings at a resolution of 1 kb on the region from 1,350,000 to 1,600,000 bp of chromosome 7.

## Availability of data and materials

The datasets and code supporting the conclusions of this article are available at https://github.com/zhengchangsulab/EnhancerSilencerPrediction and are included within the article and its additional files.

## Acknowledgements

We thank the ENCODE, 4DN, CISTROME and silencerDB databases for providing the RNA-seq data, Hi-C data, epigenetic data, and silencer related data.

## Author information

### Authors and Affiliations

Department of Bioinformatics and Genomics, the University of North Carolina at Charlotte, Charlotte, NC, 28223, USA

Sisi Yuan, Pengyu Ni & Zhengchang Su

Department of Molecular Biophysics and Biochemistry, Yale University, New Haven, CT 06520, USA Pengyu Ni

### Contributions

ZS and SY designed the study. SY performed the bioinformatic analysis with contributions from PN and ZS. SY and ZS wrote the manuscript with input from all co-authors. All authors read and approved the final manuscript.

### Corresponding author

Correspondence to Zhengchang Su.

## Ethics declarations

### Ethics approval and consent to participate

Not applicable.

### Consent for publication

Not applicable.

